# MRI-Derived Markers of Acute and Chronic Inflammatory Processes in the VTA Associated with Depression

**DOI:** 10.1101/2025.04.30.651588

**Authors:** Sarah Khalife, Steffen Bollmann, Andrew Zalesky, Lena Oestreich

## Abstract

**Background:** Depression is a leading cause of disability worldwide, with inflammation increasingly recognized as a contributing factor. Inflammatory processes can disrupt the brain’s reward circuitry, particularly the ventral tegmental area (VTA), which is central to dopamine-mediated motivation and reward. This study investigates whether MRI-derived markers sensitive to neuroinflammation and microstructure in the VTA are associated with depression diagnosis and symptom severity.

**Methods:** We analyzed diffusion weighted imaging and quantitative susceptibility mapping data from 32,495 UK Biobank participants, including 3,807 individuals with ICD-10 diagnosed depression. Metrics sensitive to neuroinflammation (free water [FW], isotropic volume fraction [ISOVF], magnetic susceptibility) and microstructure (intracellular volume fraction [ICVF], orientation dispersion index [ODI] volume) were extracted from the VTA. Group differences between the major depression group and BMI, sex, and age-matched healthy controls were assessed using ANOVAs and linear regression was used to predict acute symptom severity based on Recent Depressive Symptoms scores.

**Results:** Participants with depression diagnosis had significantly higher FW (*p* < 0.001) and ISOVF (*p* = 0.001) compared to HCs, indicating increased extracellular processes such as inflammation in the VTA. Lower ISOVF (*β* = -0.46, *p* = 0.017) and higher ICVF (*β* = 0.34, *p* = 0.007) and ODI (*β* = 0.49, *p* = 0.004) were associated with higher depression severity, independent of depressive diagnosis history.

**Conclusions:** Our findings reveal distinct patterns of VTA microstructural changes associated with depression history versus acute depressive symptom severity, suggesting different underlying pathophysiological mechanisms. Distinct patterns of neuroinflammation may differentiate acute from chronic depression, informing targeted interventions.

## Introduction

According to the World Health Organization (WHO), depression affects approximately 300 million individuals worldwide, making it a leading cause of disability (1). Depression is a complex, multifactorial disorder, with growing evidence suggesting that inflammation plays a crucial role in its pathophysiology, contributing to both its onset and progression (2). Studies report that individuals with depression often exhibit inflammatory markers and cytokines in their cerebrospinal fluid and blood, suggesting the activation of microglia and astrocytes (illustrated in Figure 1); (3). These inflammatory responses can be triggered by various environmental factors, prolonged stress, infections, neurological conditions, and autoimmune diseases, amongst others (4,5). Chronic inflammation may lead to oxidative and nitrosative stress, causing neurotoxicity and neuronal damage, both of which have been linked to depression (6).

**Figure 1.**
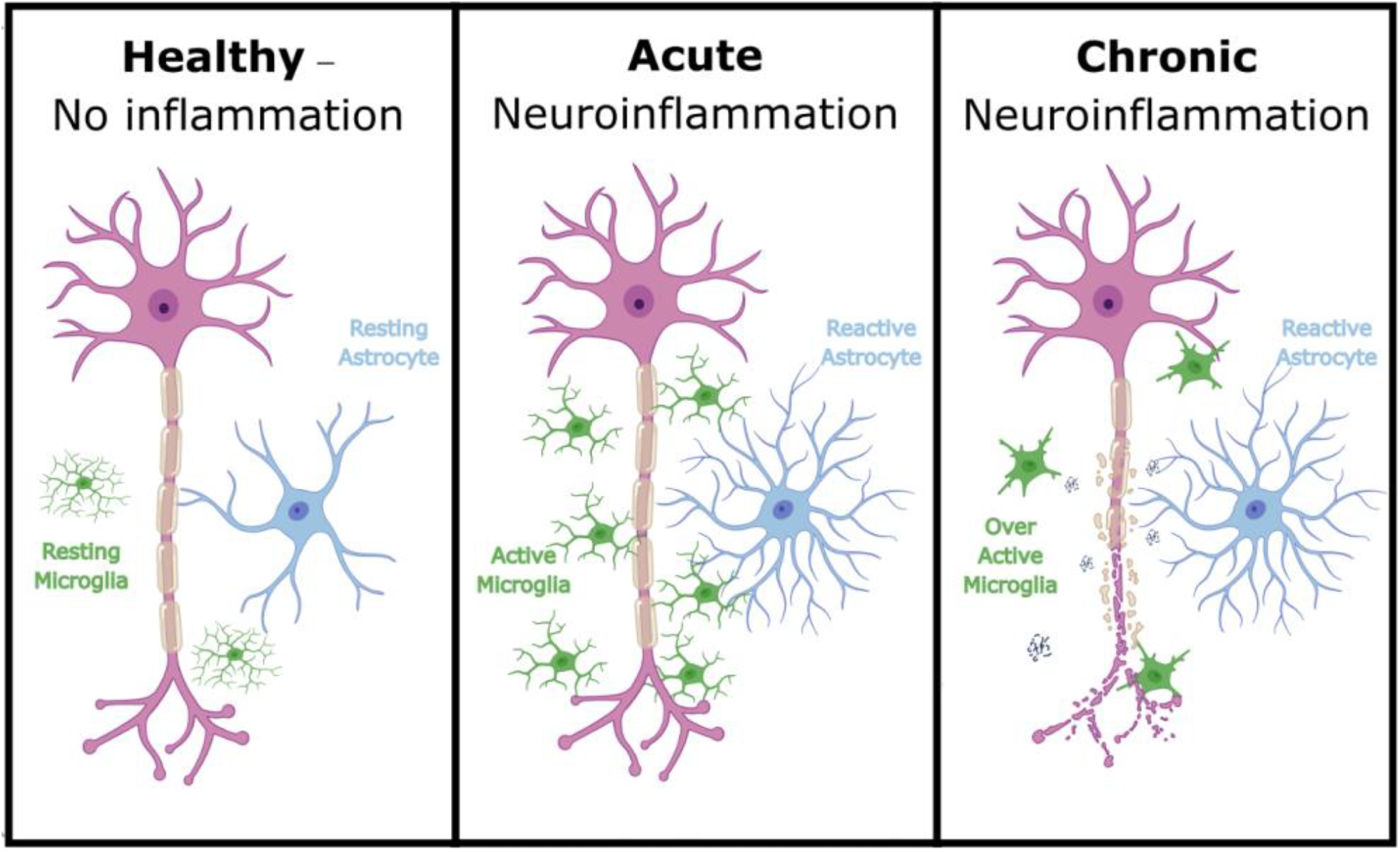
Cellular changes in response to varying degrees of inflammation. The figure illustrates the progression of neuroinflammation from a healthy state to acute and chronic inflammation. In the absence of inflammation, microglia (in green) exhibit a resting morphology with fine, branched processes, and astrocytes (in blue) maintain their characteristic star-shaped structure. During acute neuroinflammation, microglia become activated, adopting an amoeboid shape, while astrocytes display hypertrophy, indicating reactivity. In chronic neuroinflammation, sustained elevated cytokine levels cause prolonged activation in both cell types: microglia become overactive, while astrocytes remain reactive. This results in cellular toxicity and potential neurodegeneration, characterized by increased iron deposition and myelin sheath breakdown.

Inflammation disrupts the brain’s reward network, which is central to regulating motivation and pleasure (7). Anhedonia and lack of motivation, hallmark symptoms of depression, may be linked to the impact of inflammatory cytokines on mesolimbic dopamine signaling (8). Previous research suggests that inflammatory processes can disrupt the synthesis, release, and reuptake of dopamine, potentially reflecting an adaptive mechanism that prioritizes energy conservation over reward-seeking behaviors during inflammatory states (9). This notion is further supported by animal models demonstrating that inflammation reduces reward-seeking behaviors (10,11).

At the center of the reward network is the ventral tegmental area (VTA), a rich dopaminergic region that plays a critical role in regulating mood, motivation and reward (12). The VTA is highly susceptible to ischemia and inflammation, which can lead to neuronal dysfunction in this region (13). Although previous studies have reported decreased VTA activation in depression, microstructural changes in the VTA remain underexplored, likely due to its small, complex structure (14). However, recent advancements in neuroimaging, such as high-resolution parcellation atlases, now enable more precise VTA mapping (15).

Several neuroimaging studies have shown a relationship between inflammation and depression. For instance, Positron Emission Tomography (PET) studies have revealed increased radiotracer uptake by microglia in depressed subjects, indicating heightened inflammation (16). Other imaging metrics linked to inflammation, such as increased magnetic susceptibility in various brain regions measured using quantitative susceptibility mapping (QSM), have also been implicated in depression (17,18). Diffusion weighted imaging (DWI) has similarly revealed inflammation-related microstructural changes (19) and elevated free water (FW) in the extra cellular space, commonly associated with inflammatory processes (20). For instance, patients with post-stroke depression, exhibit elevated FW in reward-related regions such as the amygdala, hippocampus, and areas also associated with systemic inflammation in depression (21,22).

In this study, we utilize QSM and DWI to non-invasively examine whether putative markers sensitive to inflammation, as well as microstructural markers that may be altered as a secondary consequence of inflammation are associated with depression in the VTA. Given the established relationship between inflammation, dopaminergic dysfunction, and depression, as well as the VTA’s central role in reward circuitry, we hypothesize that individuals with a history of major depressive disorder will exhibit significantly higher levels of QSM- and DWI-derived inflammatory markers and microstructural changes in the VTA compared to matched healthy controls (HC). Based on the transient nature of inflammation, its impact on dopamine dysfunction – leading to a loss of motivation – and its effect on brain morphology, we furthermore hypothesized that acute depression severity, can be predicted by MRI markers sensitive to both inflammation and microstructural changes in the VTA irrespective of diagnostic history of depression.

## Materials and Methods

### Participants

The UK Biobank (UKB) is a large-scale, prospective epidemiological cohort study comprising approximately 500,000 participants recruited from the general UK population (23). Here, we utilized a subsample of the UKB for whom DWI, QSM, and T1-weighted MRI of brain anatomy was acquired (N=46,703). Participants with disorders known to impact brain structure, were excluded, including individuals with: (1) intellectual disabilities (2) psychotic disorders or bipolar disorder, and (3) a history of head injury or neurological conditions (**see Supplementary Table 1 for the detailed list of variables**). We opted not to systematically exclude or control for immunometabolic or inflammatory comorbidities, primarily because the UK Biobank does not provide sufficiently detailed inflammatory markers to reliably identify individuals’ inflammatory status at the time of scanning, and because excluding those with common comorbidities would reduce the generalizability of our findings to real-world clinical populations. Participants with missing data or statistical outliers (values > 3 standard deviations from the mean) were also excluded. The final sample included 32,495 participants. Ethical approval was granted by the UKB (HREC reference 11/NW/0382, application 100773) and ratified by the University of Queensland ethics committee (2023/HE000221).

### Depression Measures

Of our total sample, 3,807 individuals had a history of major depressive disorder, as determined by an International Classification of Diseases, 10th Revision (ICD-10) diagnosis of a single episode or recurrent depression (UKB field ID: 41270). To compare individuals with and without a history of depression, we matched a subset of participants on BMI, age, and sex. Descriptive statistics for both groups are presented in Table 1.

**Table 1.** Summary of demographic and MRI-derived metrics for the study sample, including total sample from the UK biobank that fit our inclusion criteria, healthy control (HC) group, and participants with a history of major depression based on ICD-10 diagnosis. Values for continuous variables are presented as means with standard deviations in parentheses. BMI = body mass index BMI, RDS = recent depressive symptoms scores, VTA = ventral tegmental area, ICVF = including intra-cellular volume fraction, ODI = orientation dispersion index, ISOVF = isotropic volume fraction, FW = free water. P-values represent group comparisons conducted using independent t-tests, with significance levels denoted as ***p < 0.001, **p < 0.01, and *p < 0.05.

| Variable | Total | HC | Depression History | p-value |
| --- | --- | --- | --- | --- |
| N | 32,495 | 3,807 | 3,807 |  |
| Female n (%) | 16,806 (51.7%) | 2,435 (64%) | 2,431 (63.9%) |  |
| Age (Years) | 63.9 (7.7) | 62.4 (7.6) | 62.5 (7.6) | 0.788 |
| BMI (kg/m<sup>2</sup>) | 26.2 (3.9) | 26.9 (4.2) | 26.9 (4.2) | 0.740 |
| RDS | 5.2 (1.7) | 5.1 (1.6) | 6.5 (2.6) | p<0.001*** |
| ***MRI Derived Metrics*** |  |  |  |  |
| ICVF | 0.533 (0.074) | 0.531 (0.073) | 0.527 (0.072) | 0.044* |
| ODI | 0.295 (0.073) | 0.294 (0.073) | 0.298 (0.073) | 0.025* |
| ISOVF | 0.374 (0.114) | 0.366 (0.115) | 0.375 (0.115) | 0.001** |
| FW | 0.456 (0.101) | 0.451 (0.102) | 0.459 (0.103) | p<0.001*** |
| Magnetic Susceptibility | 0.395 (0.072) | 0.393 (0.071) | 0.394 (0.072) | 0.457 |
| Volume Ratio | $7.025 \times 10^{-4}$ ($1.98 \times 10^{-4}$) | $7.135 \times 10^{-4}$ ($2.057 \times 10^{-4}$) | $7.076 \times 10^{-4}$ ($2.006 \times 10^{-4}$) | 0.201 |

To address the possibility that an ICD-10 diagnosis may not reflect each participant’s current depressive state—given that some diagnoses may have been established years before the imaging and individuals could now be in remission or symptom-free—we used the total score from the Recent Depressive Symptoms (RDS) questionnaire to quantify acute depressive symptoms, irrespective of depression history. Completed on the same day as the neuroimaging session, the RDS assessed four self-reported depressive symptoms over the past two weeks: 1) depressed mood, 2) unenthusiasm/disinterest, 3) tenseness/restlessness, and 4) tiredness/lethargy. Each symptom was rated on a 4-point Likert scale (1 = not at all, 4 = nearly every day), resulting in a sum score ranging from 4 to 16, where higher scores indicated more frequent and severe depressive symptoms (UKB field IDs: 2050, 2060, 2070, 2080). The reliability of the RDS has been previously cross-validated against other established depression scales including PHQ-9 (24).

### Imaging acquisition and processing

DWI, susceptibility weighted, and T1-weighted MRI images were acquired by the UKB at several sites on a 3T Siemens Skyra scanner with a 32-channel radiofrequency receive head coil, using a standardized acquisition protocol (25).

T1-weighted images were acquired using a 3D MPRAGE acquisition, with 1mm isotropic resolution. DWI images were acquired using a multi-band spin echo EPI sequence (25) at 2mm isotropic resolution. A multishell approach with two b-values (b = 1000 s/mm^2^ and b = 2000s/mm^2^, 5 non-diffusion weighted images of b = 0 s/mm^2^ and one reverse-phase encoded b = 0 s/mm^2^) were used for the acquisition and 50 diffusion encoding directions, acquired for each diffusion-weighted shell. Preprocessing of the images included gradient distortion, slice outlier, head motion and eddy currents corrections (25). FW-corrected maps were derived from the pre-processed DWI tensor maps (26). Neurite orientation dispersion and density imaging (NODDI) was modeled from DWI (27) by the UKB using the AMICO tool (28). The NODDI measures used in our study were the volume fraction of Gaussian isotropic volume fraction (ISOVF), intra- cellular volume fraction (ICVF) and orientation dispersion indices (ODI). Susceptibility-weighted imaging was used to derive QSM images which were minimally pre-processed by the UKB to remove background field artifacts and outlier brain-edge voxels with high phase variance (25). To efficiently compare QSM with other variables, we applied min-max normalization, scaling the data between 0 and 1 and centering it on the minimum value.

Neuroimaging analyses were conducted on the high-performance computer Bunya located at the University of Queensland (29), utilizing Neurodesk (30). The VTA was delineated with the Levinson-Bari Limbic Brainstem Atlas (15) by co-registering the T1 images to DWI images using Advanced Normalization Tools (ANTs) (31). Then, a T1 MNI template (32) was co-registered to the T1 image in DWI subject space. The resulting transformation matrix was then used to register the VTA to DWI subject space (illustrated in Figure 2). QSM images were similarly co-registered to DWI subject space.

**Figure 2.**
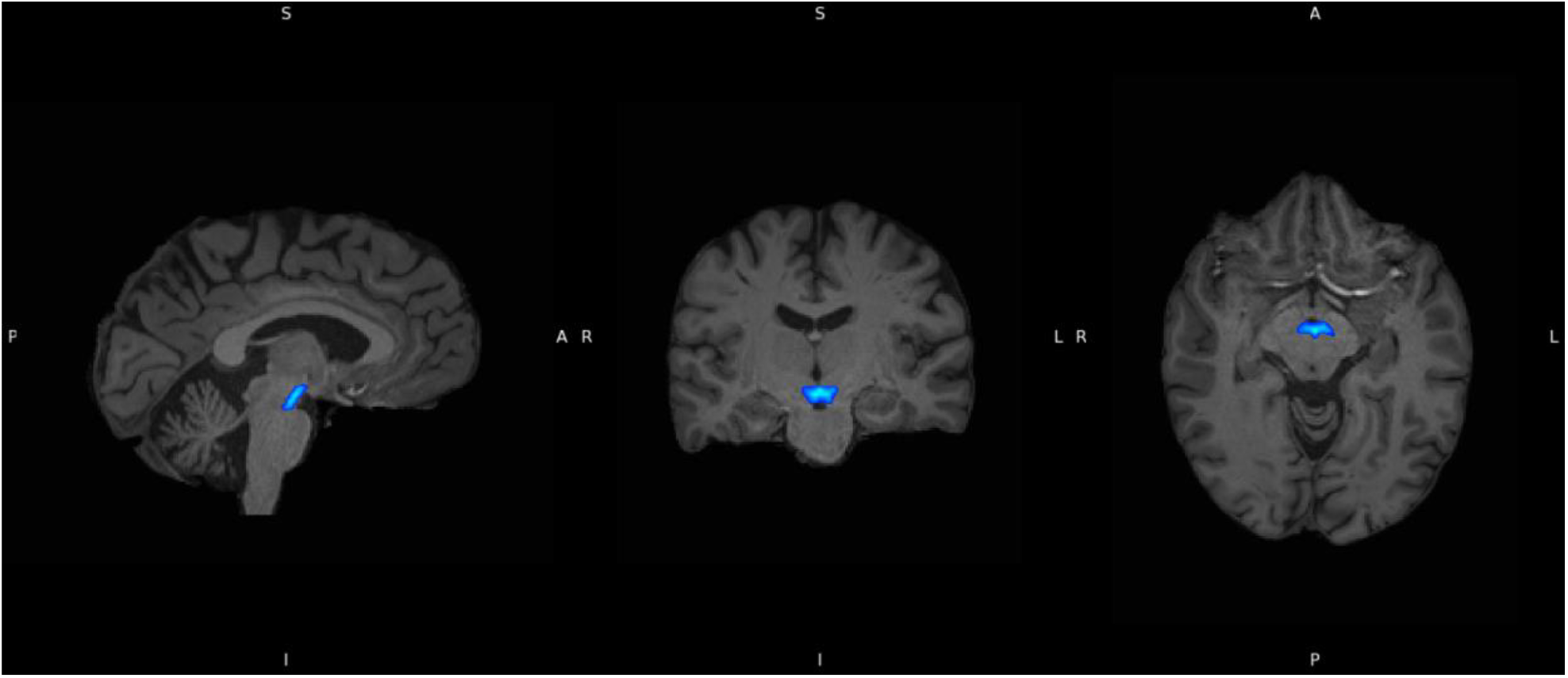
Ventral Tegmental Area (in blue), delineated using the Levinson-Bari Limbic Brainstem Atlas, displayed on co-registered subject T1 images. Orientation markers: R = Right, L = Left, A = Anterior, P = Posterior, S= Superior, I= Inferior.

Estimates of the diffusion metrics FW, ISOVF, as well as magnetic susceptibility were extracted from the VTA, as these metrics are reportedly associated with extracellular processes such as neuroinflammation (33–35). Although both FW and ISOVF measure free water in tissue, ISOVF is derived from NODDI and requires advanced diffusion protocols, while FW is simpler to acquire and thus more feasible in routine clinical settings. By testing both, we aimed to determine whether FW—a more accessible clinical measure—could reliably capture the neuroinflammatory changes indicated by ISOVF, thereby broadening clinical utility in identifying depression-related inflammation. ICVF and ODI were additionally extracted to investigate potential microstructural changes associated with acute and prolonged neuroinflammation.

Volume of the VTA was extracted and standardized using the total intracranial volume to decrease the effects of individual differences in brain size that might in turn affect the size of the VTA. Standardized VTA volume was estimated to investigate changes potentially indicative of localized neuronal depletion.

### Statistical Analysis

Group differences on demographic and clinical metrics were tested using t-tests. To investigate whether MRI-derived markers sensitive to neuroinflammation and structural changes in the VTA are associated with depression diagnosis, we conducted a series of one-way analysis of variance (ANOVA)s with *group* (depression/no depression) as between-groups factor and *VTA derived-metrics* (FW, ISOVF, QSM, ICVF, ODI, volume) as within-groups factor. Groups were matched on age, sex, and BMI. Bonferroni correction was applied to account for multiple comparisons, setting the corrected alpha level at 0.0083 (0.05/6 comparisons). Effect sizes were reported as eta squared (η^2^).

To investigate whether MRI metrics in the VTA predicted current depressive symptoms severity, we performed a linear regression analysis using MRI markers as predictors and depression severity as outcome variable. Since approximately 75% of participants had low RDS scores of 4 or 5 (see **Supplementary Figure 1**), we selected a subset of individuals to balance the distribution of RDS scores. To ensure adequate representation of individuals with high depressive symptom severity, we identified a reference group with RDS scores ≥ 13, indicating at least one depressive symptom experienced daily over the past two weeks. Given the limited number of participants in this high-severity reference group, (< 100 participants per RDS score above 13), we limited the sampling of participants with lower RDS scores to a maximum of 100 participants per RDS score.

Selection of 100 participants for each RDS score < 13 was done pseudo-randomly to minimize differences in average age and sex ratios compared to the reference group. An iterative process (with 50,000 permutations) was used to closely match the age and sex distributions of the reference group. Exact matching was deemed too restrictive, often resulting in fewer than 100 subjects per score. To address residual differences, we controlled for the potential confounders age, age squared, sex, and BMI in the regression analysis. We used standardized coefficients to allow direct comparisons of effect sizes across all predictors. Robustness was ensured by assessing multicollinearity among independent variables using the variance inflation factor (VIF) and exploring variable relationships through correlations analysis (**Supplementary Figure 2**).

## Results

### Association Between VTA-Derived MRI Markers and Depression Diagnosis

As shown in Table 1, our groups consisted of 64% females and had an average age of 62.5±7.6 years and a BMI of 26.9±4.2 kg/m^2^. There were no significant group differences in age, sex, or BMI, indicating that these demographic variables were well-matched across groups. As expected, the depression history group exhibiting notably higher RDS scores compared to the HC group (*t*(6327) = 28.3, *p* < 0.001).

ANOVAs found that the group of individuals with a history of depression exhibited significantly higher FW (*F*(1, 7612) = 12.29, *p* < 0.001, *η*^*2*^ = 0.0016) and ISOVF (*F*(1, 7612) = 10.26, *p* = 0.001, *η*^2^ = 0.0013) compared to the HCs in the VTA, suggesting increased extracellular inflammation-related processes amongst individuals with a history of depression. Higher ICVF (*F*(1, 7612) = 4.06, *p* = 0.0439, *η*^2^ = 5.34*e*^*-*04^) and ODI (*F*(1, 7612) = 4.997, *p* = 0.0254, *η*^2^ = 6.56*e*^-04^) in the depression group compared to the HC group did not survive Bonferroni correction. No significant group differences were observed for volume (*F*(1, 7612) = 1.632, *p* = 0.2, *η*^2^ = 0.0002) or magnetic susceptibility (*F*(1, 7612) = 0.554, *p* = 0.457, *η*^2^ = 7.2*e*^*-*05^). Group means and distributions are visualized in Figure 3

**Figure 3.**
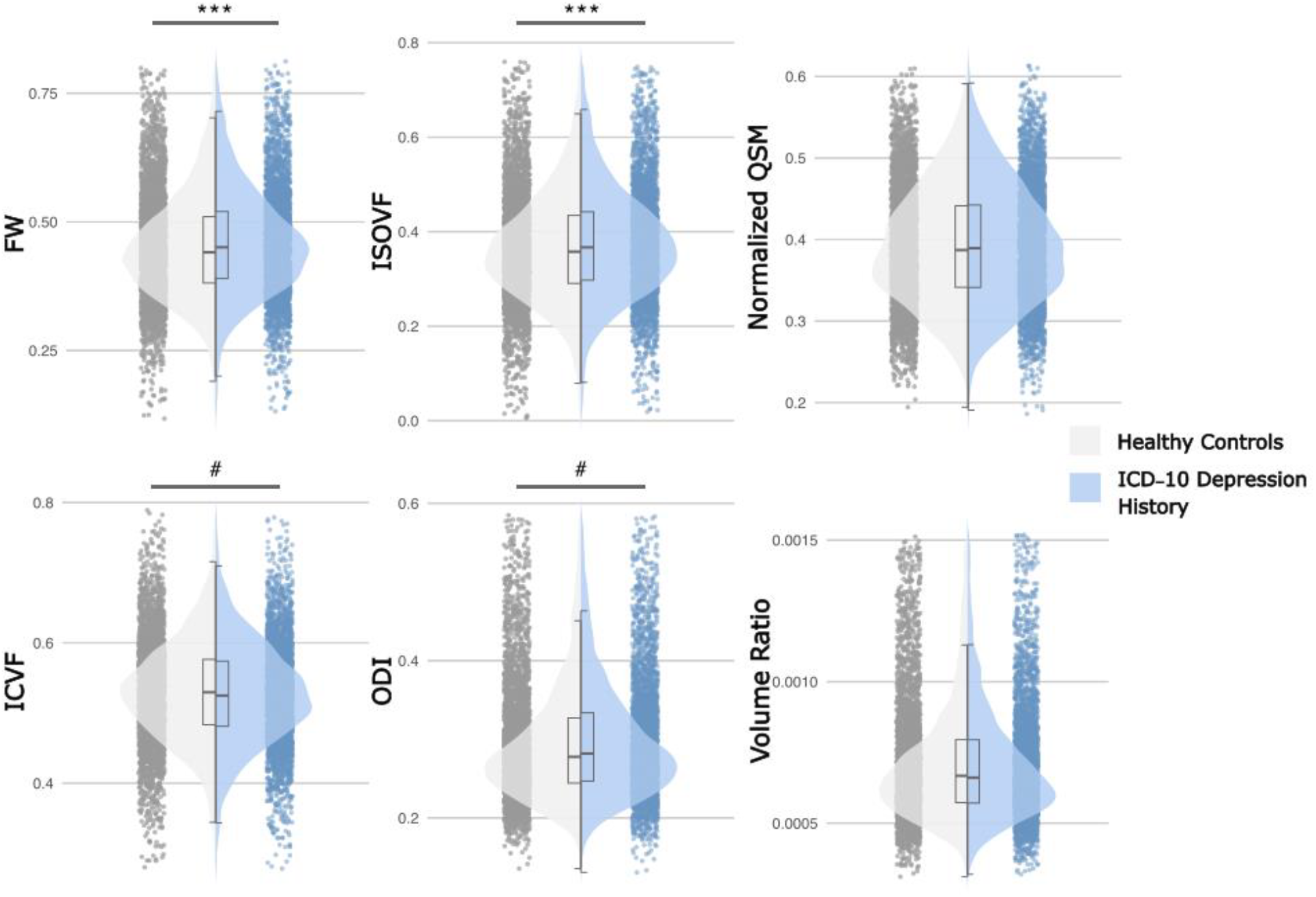
Group comparisons between healthy controls (HC) and individuals with an ICD- 10 history of depression. (*) represents significant group differences that sustained even after multiple comparisons with (***) representing a p-value ≤ 0.001 and (#) indicates significant group differences before multiple corrections. MRI metrics from the ventral tegmental area (VTA), including intra-cellular volume fraction (ICVF), orientation dispersion index (ODI), isotropic volume fraction (ISOVF), free water (FW), quantitative susceptibility mapping (QSM), and standardized volume ratio.

### VTA-Derived MRI Markers as Predictors of Acute Depression Severity

The final subsample, comprising participants selected from both the reference groups (RDS scores ≥ 13) and the best-matched groups (RDS scores < 13), consisted of a total of 1,151 participants after RDS score sampling. Detailed descriptive statistics for total subsample, as well as for the best-matched and reference groups, are presented in Table 2. The total subsample consisted of approximately 58% females, with a mean age of 60.9±7.4 years and a BMI of 27.2±4.4 kg/m^2^.

**Table 2.** Demographic and MRI-derived metrics of the total sample from the UK biobank that fit our inclusion criteria and the total subsample which consisted of individuals with varying levels of recent depressive symptom (RDS) scores and was the sum of both the best-matched (RDS < 13) and reference (RDS ≥ 13) groups. BMI = body mass index BMI, RDS = recent depressive symptoms scores, VTA = ventral tegmental area, ICVF = including intra-cellular volume fraction, ODI = orientation dispersion index, ISOVF = isotropic volume fraction, FW = free water.

| Variable | Total Subsample | Best matched<br>(RDS<13) | Reference<br>(RDS ≥ 13) |
| --- | --- | --- | --- |
| N | 1,151 | 900 | 251 |
| Female n (%) | 667 (57.9%) | 506 (56.2%) | 161 (64.1%) |
| Age (Years) | 60.9 (7.4) | 61.3 (7.5) | 59.5 (6.9) |
| BMI (kg/m<sup>2</sup>) | 27.2 (4.4) | 26.9 (4.2) | 27.9 (5) |
| RDS | 9.4 (3.5) | 8 (2.6) | 14.4 (1.2) |
| ***MRI Derived Metrics*** |  |  |  |
| ICVF | 0.529 (0.074) | 0.527 (0.074) | 0.533 (0.074) |
| ODI | 0.297 (0.077) | 0.297 (0.078) | 0.296 (0.076) |
| ISOVF | 0.371 (0.115) | 0.375 (0.115) | 0.356 (0.115) |
| FW | 0.455 (0.103) | 0.459 (0.103) | 0.441 (0.104) |
| Magnetic Susceptibility | 0.392 (0.07) | 0.391 (0.069) | 0.394 (0.073) |
| Volume Ratio | $7.128 \times 10^{-4}$ ($2.01 \times 10^{-4}$) | $7.125 \times 10^{-4}$ ($2.01 \times 10^{-4}$) | $7.139 \times 10^{-4}$ ($2.016 \times 10^{-4}$) |

When all variables in the linear regression model were included, the VIF for FW (VIF = 34.12) and ISOVF (VIF = 33.08) were substantially elevated, indicating a high degree of multicollinearity. Correlation analysis showed a strong relationship between FW and ISOVF (*r* = 0.96, *p* < 0.001), suggesting that these two variables likely reflect similar processes, allowing them to be used interchangeably. Thus, due to ease of interpretation in the context of the NODDI metrics, FW was excluded from the regression model while ISOVF was retained as a proxy measure for elevated extracellular processes linked to inflammation. After excluding FW, ISOVF VIF value decreased to 1.82 and remaining variables other than age and age squared showed VIF values below 5, suggesting that multicollinearity was adequately addressed. Refer to Supplementary Figure 2 for more variable correlations.

The overall linear regression model was significant (*F*(9, 1141) = 9.673, *p* < 0.001) and explained 6.35% of the variance in depression severity (*R*^2^ = 0.0709, *R*^2^ adjusted = 0.0635). Model fit was assessed in Figure 4(A), showing a moderate overall fit with a tendency to overestimate predicted values near the center of the distribution, whereas Figure 4(B) displays standardized beta coefficients for each predictor, illustrating their relative impact on depression severity and highlighting which factors most strongly influence symptoms. Both ICVF (*β* = 0.34, *p* = 0.007, 95% CI [0.93, 0.58]) and ODI (*β* = 0.49, *p* = 0.004, 95% CI [0.15, 0.83]) were positively associated with depressive symptom severity, indicative of an increase in microstructural changes with higher depression severity. Elevated ODI values suggest increased microstructural changes in the extracellular space, while higher ICVF values indicate greater neurite density associated with worsening depression symptoms. ISOVF showed a negative association with depression severity (*β* = -0.32, *p* = 0.017, 95% CI [-0.59, -0.06]), indicating that lo in the free-water component was linked to greater symptom severity. In contrast, standardized volume and magnetic susceptibility were not significantly associated with depressive symptoms.

**Figure 4.**
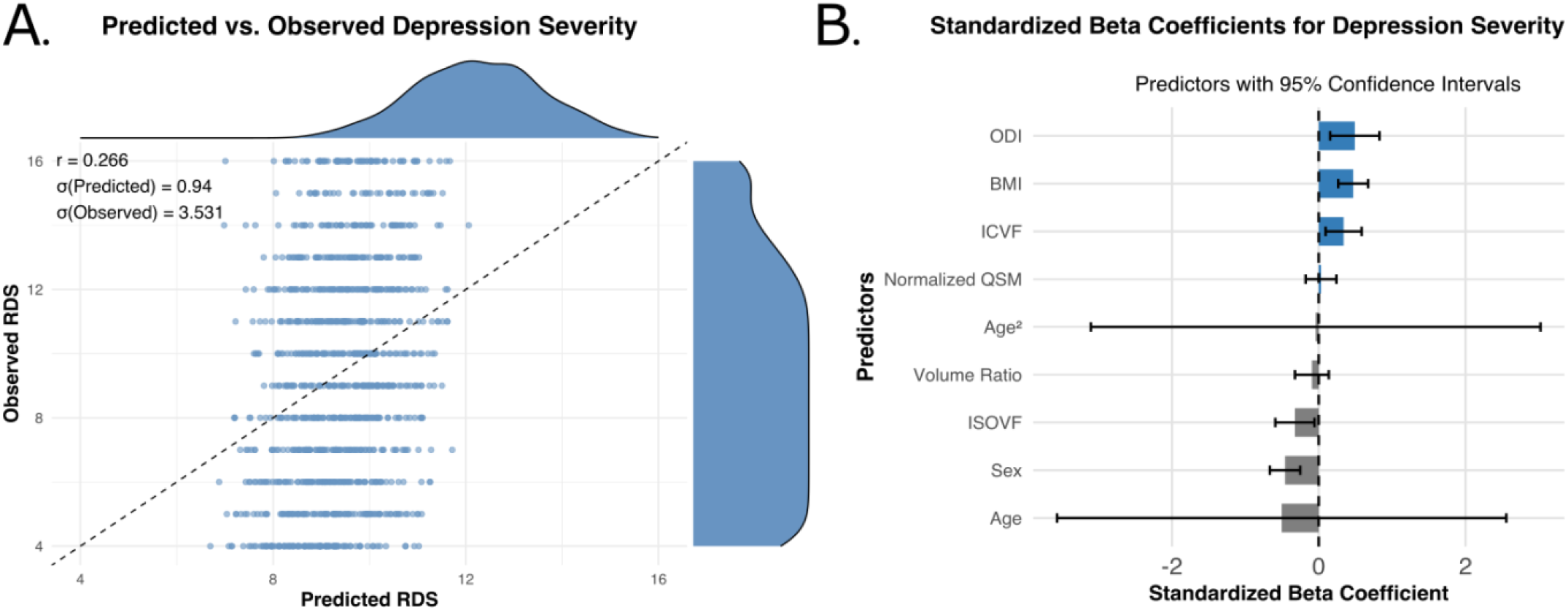
(A) Scatter plot depicting the relationship between predicted and observed depression severity scores (RDS) derived from the linear regression model. Each point represents an individual participant, with the dashed identity line (y = x) included for reference. The marginal density plots illustrate the distribution of predicted and observed scores. The Pearson correlation coefficient (r), along with the standard deviations of the predicted and observed RDS values, is annotated. (B) Standardized beta coefficients from the regression model predicting depression severity. Bars represent the effect size of each predictor, with 95% confidence intervals indicated by error bars. Positive predictors (e.g., ODI, BMI, ICVF) are shown in blue, while negative predictors are shown in grey. The dashed horizontal line at zero represents the null effect. Variables are ordered by effect size for clarity.

Sex was a significant predictor of depression severity, whereby males exhibited lower depressive symptom severity compared to females (*β* = -0.46, *p* < 0.001, 95% CI [- 0.67, -0.25]). Additionally, BMI was positively associated with depressive severity (*β* = 0.47, *p* < 0.001, 95% CI [0.26, 0.67]), suggesting that higher BMI may be linked to increased severity of depressive symptoms. The covariates age and age squared were not significantly associated with depressive symptom severity.

## Discussion

This study provides novel insights into the neurobiological correlates of depression, revealing a complex relationship between structural and inflammatory markers in the VTA and depressive symptomatology. Our findings demonstrate that individuals with a diagnosis of depression exhibited significantly higher extracellular free-water in the VTA compared to matched HC, suggesting extra-neuronal pathology potentially linked to inflammatory processes in the major depression group. While these inflammatory markers were elevated, no significant structural alterations in the VTA or difference in iron deposition or demyelination indicators, as measured by QSM, were observed between groups with and without a history of depression. Notably, MRI markers sensitive to inflammation and structural changes in the VTA predicted acute depressive symptom severity, independent of prior history of depression diagnosis. Specifically, ICVF, ODI and ISOVF in the VTA emerged as independent predictors of acute depression severity, whereby ICVF and ODI were more elevated, while ISOVF was lower with higher depression severity. In contrast, magnetics susceptibility did not predict acute depression severity, aligning with our finding of no significant differences between the HC and depression history groups, thus supporting an absence of chronic inflammation effects like cell damage or iron deposition.

ISOVF and FW are MRI markers sensitive to neuroinflammation and edema (26,27). While both ISOVF and FW are diffusion-based measures that quantify freely diffusing water content in brain tissue through distinct modeling approaches (three-compartment NODDI versus two-compartment tensor model respectively), both measures can reflect aspects of inflammatory processes, which are typically characterized by increased extracellular water content arising from vasogenic edema and associated cellular responses (26). As such, elevated ISOVF and FW in individuals with a history of depression aligns with our prediction that major depression is associated with elevated inflammation markers in the VTA. In contrast, while FW and ISOVF were higher in individuals with major depression, we observed that acute depression severity in the general population was associated with lower ISOVF. This dissociation suggests these two states of depression - historical diagnosis of major depression versus acute depressive symptoms across the general population - may reflect distinct biological processes and inflammatory states. Yi et al. (36) highlighted important distinctions between acute and chronic inflammation, showing that acute neuroinflammation is marked by increased hindered water diffusion due to increases in the number of activated hyper-ramified microglia, while chronic inflammation is marked by reduced hindered diffusion due to overactivation of microglia which increases their density. Thus, our findings of lower ISOVF might reflect acute inflammation processes, while the elevated ISOVF and FW might be indicative of chronic inflammatory processes in major depression.

Increased FW is often associated with psychiatric symptom severity and neuroinflammation in psychiatric and neurological disorders (20,37–39). However, evidence on FW in depression remains mixed. For instance, elevated FW has been reported in post-stroke depression patients in reward-related regions (22), suggesting that brain areas critical for reward processing are especially susceptible to neuroinflammation. Conversely, Bergamino et al. (37) found no differences in FW between HC and unmedicated MDD patients. Consistent with our findings, Al Thubaity et al. (40) observed a general reduction in ISOVF levels in the grey matter of depressed individuals compared to HC. Specifically, ISOVF reductions were noted in the anterior cingulate cortex (ACC) of depressed patients with low peripheral inflammation, although this difference was not statistically significant across the entire depression group. The ACC, like the VTA, has shown decreased fMRI activation and increased microglial expression in PET imaging (41). However, Thubaity et al. (40) found no significant ISOVF differences in the prefrontal regions or insular cortex when comparing high/low inflammation subgroups or when comparing the overall depression group to controls. These findings suggest that FW and ISOVF changes may reflect distinct, localized processes, potentially influenced by acute or chronic inflammation associated with depression. Given the reward network’s sensitivity to neuroinflammatory processes and its central role in dopamine signaling (9- 11), these regions might exhibit differential responses to such changes with prolonged depressive episodes possibly impacting the VTA’s structural integrity in a manner distinct from acute depressive symptoms.

Our results indicate that the VTA is also susceptible to microstructural aberrations possibly linked to inflammation, in depression. While ICVF and ODI differences did not survive Bonferroni correction in group comparisons, our regression analysis identified ICVF and ODI as an independent predictor of depression severity. Elevated ICVF, which is associated with intracellular water diffusion, is typically interpreted as increased neurite density, often suggesting heightened neuronal density and neurogenesis (42). However, recent studies suggest links between water diffusion and glial cell swelling and microglial reactivity - processes strongly associated with depression (43). Similar increases in ICVF have been reported in the striatum following interferon-alpha treatment, which triggers inflammation (44). Although fatigue (a symptom of depression) increased after interferon- alpha administration, no significant rise in depression severity was observed, suggesting that inflammation affects reward circuitry even though it did not directly amplify mood symptoms (44). This may explain the vulnerability of the VTA and striatum to inflammation in depression, which aligns with our findings of increased ICVF in the VTA, a structure closely linked to the brain’s reward system, reinforcing its vulnerability to inflammatory processes and depression. Higher ODI levels in the VTA may reflect microstructural processes in the extracellular space, where microglia modulate immune defense. Yi et al. (36) demonstrated that increased ODI correlates with microglial density, further suggesting that microglial activation could be driving these microstructural changes. Similarly, Ota et al. (45) reported reduced ODI in the thalamus of MDD individuals, and post-mortem studies found decreased astrocytic markers in the thalamus of deceased MDD individuals, lending additional support for the involvement of glial cells in depression (46).

Our study found no significant differences in VTA volume between depressed and healthy individuals. This contrasts with findings by Morris et al. (47), who reported increased VTA volume linked to alterations in neuromelanin and dopamine activity in individuals with mood disorders. The absence of volumetric differences in our study suggests that the observed microstructural changes, such as higher ICVF and ODI, may occur independently of gross volumetric changes.

Even though QSM is sensitive to the effects of prolonged inflammation, our study found no significant differences in magnetic susceptibility between individuals with major depression and controls in the VTA, nor was it a predictor for acute depressive severity. This suggests that processes like iron deposition, demyelination, or cell death, typically associated with chronic inflammation, may not significantly impact VTA structure as hypothesized. Yet, Yao et al. (18) reported higher susceptibility values correlating with current depression severity in the putamen and thalamus. Duan et al. (17) found that magnetic susceptibility in the thalamus, hippocampus, and putamen correlated with depression duration rather than current symptom severity, suggesting that microstructural changes accumulate over time. The discrepancy between our VTA findings and previous work in other regions might reflect the distinct cellular architecture and inflammatory vulnerability of different brain areas. The VTA, primarily composed of dopaminergic neurons and their supporting glial cells, may respond differently to inflammatory processes compared to regions like the putamen and thalamus, which have different cellular compositions and metabolic demands. While Yao et al. (18) and Duan et al. (17) demonstrated susceptibility changes in striatal and limbic regions, our findings suggest the VTA may be more resilient to chronic inflammatory processes that lead to iron accumulation or cellular damage. Instead, the VTA appears to show more dynamic changes in extracellular space (as indicated by FW measures) that might reflect acute rather than chronic inflammatory states.

While statistically significant, our findings had small effect sizes, reflecting the multifactorial nature of depression. Depression is influenced by numerous factors, including sex, lifestyle factors, genetic predisposition, and metabolic variables such as BMI (48), which explained a substantial portion of the variance in our models. Our findings showed that females were significantly more likely to have higher depressive symptoms, consistent with previous literature indicating a higher prevalence of depression in females, potentially due to factors such as reporting tendencies and hormonal influences (49). BMI was also a positive predictor of depression severity, aligning with studies associating obesity with elevated cytokine levels, reduced physical activity, and heightened depression risk (50,51). These results underscore the complex interplay between neuroinflammation, lifestyle factors, and depressive symptoms, suggesting that future studies should consider these factors in more detail to fully understand the multifactorial nature of depression.

One important limitation of our study is the cross-sectional design, which limits our ability to establish causal relationships between neuroinflammation, VTA microstructural changes, and depression severity. Future, longitudinal studies are essential to disentangle these relationships. Additionally, while a significant portion of our subjects experienced multiple major depressive episodes between their initial ICD-10 depression diagnosis and the time of scanning, the average time since their first diagnosis was approximately 27±11.15 years prior to the scan. Future longitudinal studies should explore the relationship between recurrent depressive episodes and the interplay of chronic and acute depression on brain structure. Furthermore, while DWI and QSM provide valuable insights into brain structure and inflammation, they are indirect measures. Integrating these methods with more direct biomarkers, such as cerebrospinal fluid cytokine levels or PET imaging, would provide stronger evidence of neuroinflammation’s role in depression.

In conclusion, our findings reveal distinct patterns of microstructural changes in the VTA associated with both major depression history and current depressive symptom severity, suggesting different underlying pathophysiological mechanisms. The elevated free-water markers observed in individuals with major depression history, coupled with the relationship between NODDI metrics and current symptom severity, point to dynamic inflammatory processes that may differ between acute and chronic states of depression. While our results support the role of neuroinflammation in depression, the absence of QSM changes suggests that the VTA may be more resilient to chronic inflammatory processes compared to other brain regions. These findings could ultimately inform more targeted therapeutic approaches that consider the distinct biological processes underlying depression.

## Acknowledgements and Disclosures

LO was supported by a National Health and Medical Research Council (NHMRC) Investigator grant (2007718), a strategic award from the School of Psychology, and a start-up fund from the Australian Institute for Bioengineering and Nanotechnolgy at the University of Queensland.

This work has been previously presented as a poster abstract for the Organization of Human Brain Mapping Annual Conference 2024, Seoul, South Korea, 25th June, 2024.

Authors report no biomedical financial interests or conflicts of interest.

## Data and code availability

All the neuroimaging pre-processing and analyses conducted in this study involved the use of publicly available toolboxes and resources. Specifically, the following software tools were employed: FreeSurfer (v7.4.1) [52,53], FSL (v6.0.7.4) [54,55], FSLeyes (v1.10.4) [56], MRtrix3 (v3.0.3) [57], and ANTs (v2.3.5) [31]. For statistical analysis, both ANOVAs and regression analyses were run through R and RStudio [58], using *dplyr, ggplot2, ggpubr, skimr, plotly, htmlwidgets, MatchIt, gghalves, psych, GGally, tidyverse, car, kableExtra, officer, tcltk*, and *scales* packages and functions [59-74]. Code and materials for reproducing the analyses and figures in this study are available at https://github.com/SarahKhalife/VTA_UKB_Inflammation_Paper. Due to ethical considerations, raw data from the UK Biobank cannot be shared directly.

