## Supplementary Material for "MRI-Derived Markers of Acute and Chronic Inflammatory Processes in the VTA Associated with Depression"

Supplementary Table 1
Supplementary Figure 1
Supplementary Figure 2

#### Supplementary Table 1

List of covariates and IDs we included

| Category | Field ID | Sub field | Description |
|---|---|---|---|
| 2002 | 41270 | C70 | Malignant neoplasm of meninges |
| | | C700 | Cerebral meninges |
| | | C701 | Spinal meninges |
| | | C709 | Meninges, unspecified |
| | | C71 | Malignant neoplasm of brain |
| | | C710 | Cerebrum, except lobes and ventricles |
| | | C711 | Frontal lobe |
| | | C712 | Temporal lobe |
| | | C713 | Parietal lobe |
| | | C714 | Occipital lobe |
| | | C715 | Cerebral ventricle |
| | | C716 | Cerebellum |
| | | C717 | Brain stem |
| | | C718 | Overlapping lesion of brain |
| | | C719 | Brain, unspecified |
| | | F00 | Dementia in Alzheimer's disease |
| | | F000 | Dementia in Alzheimer's disease with early onset |
| | | F001 | Dementia in Alzheimer's disease with late onset |
| | | F002 | Dementia in Alzheimer's disease, atypical or mixed type |
| | | F009 | Dementia in Alzheimer's disease, unspecified |
| | | F01 | Vascular dementia |
| | | F010 | Vascular dementia of acute onset |
| | | F011 | Multi-infarct dementia |
| | | F012 | Subcortical vascular dementia |
| | | F013 | Mixed cortical and subcortical vascular dementia |

| | | | |
|---|---|---|---|
| | | F018 | Other vascular dementia |
| | | F019 | Vascular dementia, unspecified |
| | | F02 | Dementia in other diseases classified elsewhere |
| | | F020 | Dementia in Pick's disease |
| | | F021 | Dementia in Creutzfeldt-Jakob disease |
| | | F022 | Dementia in Huntington's disease |
| | | F023 | Dementia in Parkinson's disease |
| | | F024 | Dementia in human immunodeficiency virus [HIV] disease |
| | | F028 | Dementia in other specified diseases classified elsewhere |
| | | F03 | Unspecified dementia |
| | | F04 | Organic amnesic syndrome, not induced by alcohol and other psychoactive substances |
| | | F05 | Delirium, not induced by alcohol and other psychoactive substances |
| | | F06 | Other mental disorders due to brain damage and dysfunction and to physical disease |
| | | F07 | Personality and behavioural disorders due to brain disease, damage and dysfunction |
| | | F09 | Unspecified organic or symptomatic mental disorder |
| | | F20 | Schizophrenia |
| | | F200 | Paranoid schizophrenia |
| | | F201 | Hebephrenic schizophrenia |
| | | F202 | Catatonic schizophrenia |
| | | F203 | Undifferentiated schizophrenia |
| | | F204 | Postschizophrenic depression |
| | | F205 | Residual schizophrenia |
| | | F206 | Simple schizophrenia |
| | | F208 | Other schizophrenia |
| | | F209 | Schizophrenia, unspecified |
| | | F21 | Schizotypal disorder |

| | | | |
|---|---|---|---|
| | | F22 | Persistent delusional disorders |
| | | F220 | Delusional disorder |
| | | F228 | Other persistent delusional disorders |
| | | F229 | Persistent delusional disorder, unspecified |
| | | F23 | Acute and transient psychotic disorders |
| | | F230 | Acute polymorphic psychotic disorder without symptoms of schizophrenia |
| | | F231 | Acute polymorphic psychotic disorder with symptoms of schizophrenia |
| | | F232 | Acute schizophrenia-like psychotic disorder |
| | | F233 | Other acute predominantly delusional psychotic disorders |
| | | F238 | Other acute and transient psychotic disorders |
| | | F289 | Acute and transient psychotic disorder, unspecified |
| | | F24 | Induced delusional disorder |
| | | F25 | Schizoaffective disorders |
| | | F250 | Schizoaffective disorder, manic type |
| | | F251 | Schizoaffective disorder, depressive type |
| | | F252 | Schizoaffective disorder, mixed type |
| | | F258 | Other schizoaffective disorders |
| | | F259 | Schizoaffective disorder, unspecified |
| | | F28 | Other nonorganic psychotic disorders |
| | | F29 | Unspecified nonorganic psychosis |
| | | F30 | Manic episode |
| | | F300 | Hypomania |
| | | F301 | Mania without psychotic symptoms |
| | | F302 | Mania with psychotic symptoms |
| | | F308 | Other manic episodes |
| | | F309 | Manic episode, unspecified |
| | | F31 | Bipolar affective disorder |
| | | F310 | Bipolar affective disorder, current episode hypomanic |

| | | | |
|---|---|---|---|
| | | F311 | Bipolar affective disorder, current episode manic without psychotic symptoms |
| | | F312 | Bipolar affective disorder, current episode manic with psychotic symptoms |
| | | F313 | Bipolar affective disorder, current episode mild or moderate depression |
| | | F314 | Bipolar affective disorder, current episode severe depression without psychotic symptoms |
| | | F315 | Bipolar affective disorder, current episode severe depression with psychotic symptoms |
| | | F316 | Bipolar affective disorder, current episode mixed |
| | | F317 | Bipolar affective disorder, currently in remission |
| | | F318 | Other bipolar affective disorders |
| | | F319 | Bipolar affective disorder, unspecified |
| | | F70 | Mild mental retardation |
| | | F700 | Mild mental retardation (With the statement of no, or minimal, impairment of behaviour) |
| | | F701 | Mild mental retardation (Significant impairment of behaviour requiring attention or treatment) |
| | | F708 | Mild mental retardation (Other impairments of behaviour) |
| | | F709 | Mild mental retardation (Without mention of impairment of behaviour) |
| | | F710 | Moderate mental retardation (With the statement of no, or minimal, impairment of behaviour) |
| | | F711 | Moderate mental retardation (Significant impairment of behaviour requiring attention or treatment) |
| | | F719 | Moderate mental retardation (Without mention of impairment of behaviour) |
| | | F729 | Severe mental retardation (Without mention of impairment of behaviour) |
| | | F780 | Other mental retardation (With the statement of no, or minimal, impairment of behaviour) |

| | | | |
|---|---|---|---|
| | | F789 | Other mental retardation (Without mention of impairment of behaviour) |
| | | F790 | Unspecified mental retardation (With the statement of no, or minimal, impairment of behaviour) |
| | | F798 | Unspecified mental retardation (Other impairments of behaviour) |
| | | F799 | Unspecified mental retardation (Without mention of impairment of behaviour) |
| | | G10 | Huntington's disease |
| | | G20 | Parkinson's disease |
| | | G21 | Secondary Parkinsonism |
| | | G210 | Malignant neuroleptic syndrome |
| | | G211 | Other drug-induced secondary Parkinsonism |
| | | G212 | Secondary Parkinsonism due to other external agents |
| | | G213 | Postencephalitic Parkinsonism |
| | | G214 | Vascular parkinsonism |
| | | G218 | Other secondary Parkinsonism |
| | | G219 | Secondary Parkinsonism, unspecified |
| | | G22 | Parkinsonism in diseases classified elsewhere |
| | | G23 | Other degenerative diseases of basal ganglia |
| | | G230 | Hallervorden-Spatz disease |
| | | G231 | Progressive supranuclear ophthalmoplegia[Steele-Richardson-Olszewski] |
| | | G232 | Striatonigral degeneration |
| | | G233 | Multiple system atrophy, cerebellar type |
| | | G238 | Other specified degenerative diseases of basal ganglia |
| | | G239 | Degenerative disease of basal ganglia, unspecified |
| | | G30 | Alzheimer's disease |
| | | G300 | Alzheimer's disease with early onset |
| | | G301 | Alzheimer's disease with late onset |
| | | G308 | Other Alzheimer's disease |

| | | | |
|---|---|---|---|
| | | G309 | Alzheimer's disease, unspecified |
| | | G31 | Other degenerative diseases of nervous system, not elsewhere classified |
| | | G310 | Circumscribed brain atrophy |
| | | G311 | Senile degeneration of brain, not elsewhere classified |
| | | G312 | Degeneration of nervous system due to alcohol |
| | | G318 | Other specified degenerative diseases of nervous system |
| | | G319 | Degenerative disease of nervous system, unspecified |
| | | G320 | Subacute combined degeneration of spinal cord in diseases classified elsewhere |
| | | G328 | Other specified degenerative disorders of nervous system in diseases classified elsewhere |
| | | G35 | Multiple sclerosis |
| | | G36 | Other acute disseminated demyelination |
| | | G360 | Neuromyelitis optica [Devic] |
| | | G368 | Other specified acute disseminated demyelination |
| | | G369 | Acute disseminated demyelination, unspecified |
| | | G370 | Diffuse sclerosis |
| | | G371 | Central demyelination of corpus callosum |
| | | G372 | Central pontine myelinolysis |
| | | G374 | Subacute necrotising myelitis |
| | | G378 | Other specified demyelinating diseases of central nervous system |
| | | G379 | Demyelinating disease of central nervous system, unspecified |
| | | G90 | Disorders of autonomic nervous system |
| | | G900 | Idiopathic peripheral autonomic neuropathy |
| | | G901 | Familial dysautonomia [Riley-Day] |
| | | G902 | Horner's syndrome |
| | | G903 | Multisystem degeneration |
| | | G904 | Autonomic dysreflexia |

| | | | |
|---|---|---|---|
| | | G908 | Other disorders of autonomic nervous system |
| | | G909 | Disorder of autonomic nervous system, unspecified |
| | | G92 | Toxic encephalopathy |
| | | G930 | Cerebral cysts |
| | | G931 | Anoxic brain damage, not elsewhere classified |
| | | G932 | Benign intracranial hypertension |
| | | G933 | Postviral fatigue syndrome |
| | | G934 | Encephalopathy, unspecified |
| | | G935 | Compression of brain |
| | | G936 | Cerebral oedema |
| | | G937 | Reye's syndrome |
| | | G938 | Other specified disorders of brain |
| | | G939 | Disorder of brain, unspecified |
| | | G94 | Other disorders of brain in diseases classified elsewhere |
| | | G940 | Hydrocephalus in infectious and parasitic diseases classified elsewhere |
| | | G941 | Hydrocephalus in neoplastic disease |
| | | G942 | Hydrocephalus in other diseases classified elsewhere |
| | | G948 | Other specified disorders of brain in diseases classified elsewhere |
| | | Q01 | Encephalocele |
| | | Q02 | Microcephaly |
| | | Q03 | Congenital hydrocephalus |
| | | Q030 | Malformations of aqueduct of Sylvius |
| | | Q031 | Atresia of foramina of Magendie and Luschka |
| | | Q038 | Other congenital hydrocephalus |
| | | Q039 | Congenital hydrocephalus, unspecified |
| | | Q04 | Other congenital malformations of brain |
| | | Q040 | Congenital malformations of corpus callosum |
| | | Q043 | Other reduction deformities of brain |

| | | | |
|---|---|---|---|
| | | Q044 | Septo-optic dysplasia |
| | | Q046 | Congenital cerebral cysts |
| | | Q048 | Other specified congenital malformations of brain |
| | | Q049 | Congenital malformation of brain, unspecified |
| | | S02 | Fracture of skull and facial bones |
| | | S020 | Fracture of vault of skull |
| | | S021 | Fracture of base of skull |
| | | S022 | Fracture of nasal bones |
| | | S023 | Fracture of orbital floor |
| | | S024 | Fracture of malar and maxillary bones |
| | | S027 | Multiple fractures involving skull and facial bones |
| | | S028 | Fractures of other skull and facial bones |
| | | S029 | Fracture of skull and facial bones, part unspecified |
| | | S060 | Intracranial injury |
| | | S061 | Traumatic cerebral oedema |
| | | S062 | Diffuse brain injury |
| | | S063 | Focal brain injury |
| | | S064 | Epidural haemorrhage |
| | | S065 | Traumatic subdural haemorrhage |
| | | S066 | Traumatic subarachnoid haemorrhage |
| | | S067 | Intracranial injury with prolonged coma |
| | | S068 | Other intracranial injuries |
| | | S069 | Intracranial injury, unspecified |
| | | S07 | Crushing injury of head |

#### Supplementary Figure 1

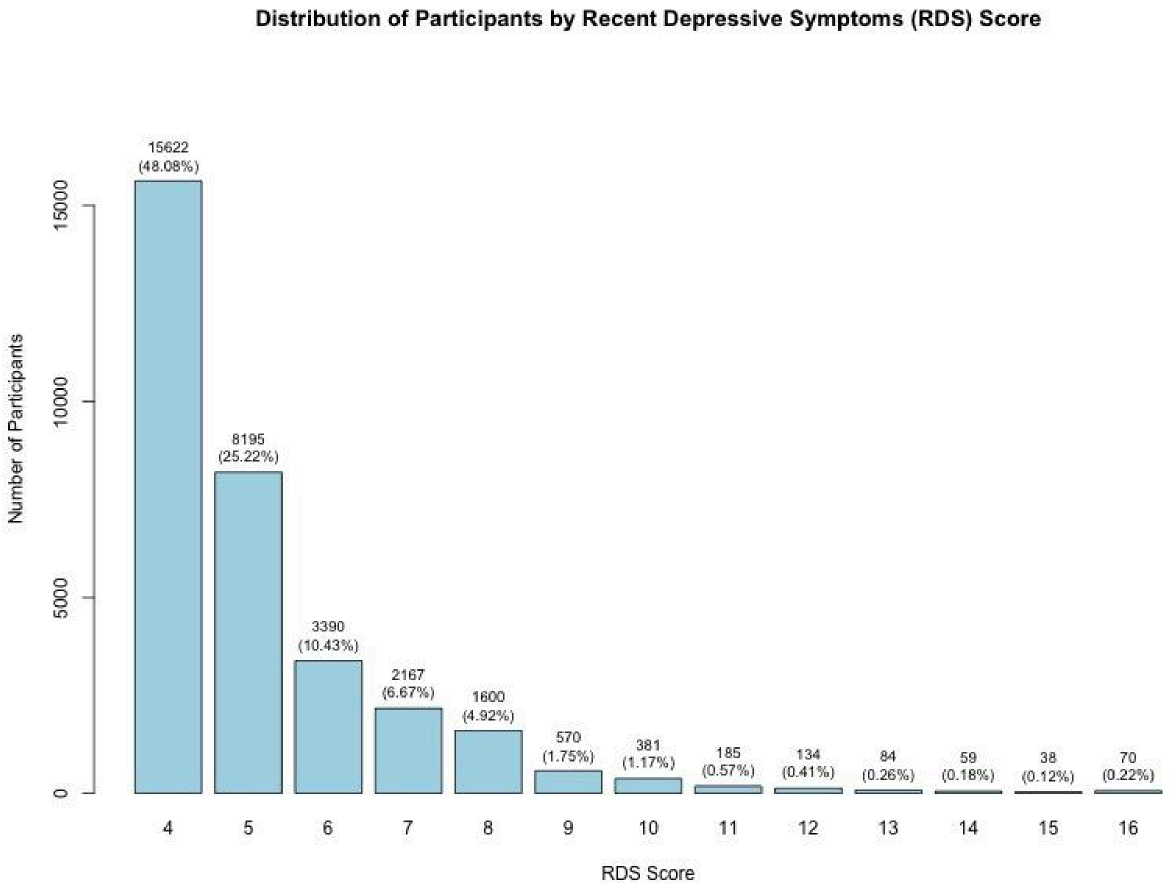


**Supplementary Figure 1.** Histogram displaying the distribution of participants across each Recent Depressive Symptoms (RDS) score. Bars represent the frequencies of participants for each RDS score, with percentages indicated above each bar.

### Supplementary Figure 2

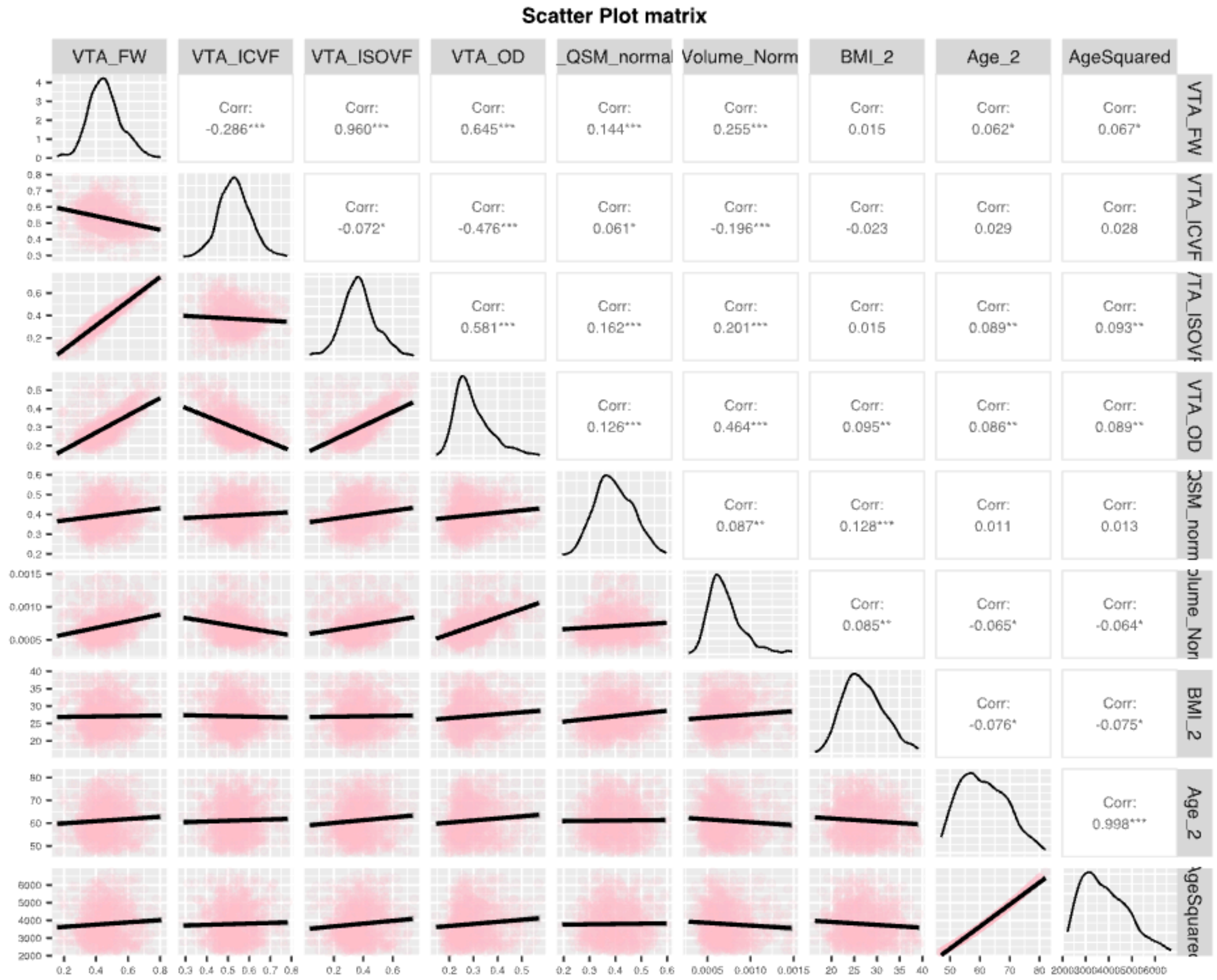


**Supplementary Figure 2.** Scatter plot and correlation matrix depicting pairwise relationships between VTA metrics and related variables. Each cell includes scatter plots showing individual data points, with correlation coefficients indicating the strength and direction of associations across variables.
